# Layer Dependence of Monocular and Binocular Responses in Human Ocular Dominance Columns at 7T using VASO and BOLD

**DOI:** 10.1101/2023.04.06.535924

**Authors:** Atena Akbari, Joseph S Gati, Peter Zeman, Brett Liem, Ravi S Menon

## Abstract

The neurons located in the striate cortex (V1) preferentially respond to the input from one eye or another, forming a fingerprint-like pattern of ocular dominance columns (ODCs). At this mesoscopic scale, accessible by ultra-high field fMRI, V1 is supplied/drained by a network of surface (pial) vessels that connect to descending/ascending tangential vessels that penetrate the cortex and supply/drain a capillary bed whose density is also layer dependent. In this study, we measured the layer dependence of monocular and binocular responses of ocular dominance columns in V1 at 7T using Blood Oxygenation Level Dependent (BOLD) and VAscular Space Occupancy (VASO) contrasts. Our results indicate that the microvascular blood volume changes that give rise to VASO are well confined to the site of neural activity across the layers of the cortex and between the columns. Pial veins dominate the BOLD response and mix the signal between columns. When the GRE BOLD response was examined in only the VASO specific voxels (thus eliminating the pial vein signal), the laminar profile was very similar to VASO, however, the columnar response was still blurred. Caution needs to be exercised in the interpretation of signal changes in BOLD at the mesoscale both in terms of feedforward/feedback effects and inhibitory and excitatory effects.

**Highlights:**

- VASO produced laminar profiles that were consistent with the known layer-dependent neuronal response to monocular and binocular stimulations.
- VASO better differentiated the response between columns belonging to the left and right eyes.
- GRE BOLD signal spatial specificity was poor in both laminar and columnar directions, however, when the pial veins were suppressed, the laminar BOLD signal was very similar to the VASO signal.
- Caution needs to be exercised when interpreting cognitive neuroscience BOLD studies at the mesoscale level due to the confounding effects of pial and sub-pial veins and venules.

## 1 Introduction

The primate primary visual cortex (V1) is divided into six histologically defined layers, with each layer having a distinct cellular and vascular composition (H.M. Duvernoy et al., 1981). The pathways of the neuronal signaling *within* each layer (lateral connections), *across* cortical layers of a brain region (intralaminar), and *between* the cortical regions (cortico-cortical) have been obtained from invasive studies on laboratory animal models (Douglas and Martin, 2004; Felleman and Van, 1991; Salin and Bullier, 1995). According to these studies of visual cortical hierarchy, thalamic input from the lateral geniculate nucleus (LGN) arrives in layers IV and VI of V1 (feedforward connections), and the feedback from higher order visual areas arrives in layers I and V of V1 (Henderickson et al., 1978; Hubel and Wiesel, 1968). In addition to this laminar organization, neurons that share the same property such as eye preference or orientation selectivity, form functional units called cortical columns that extend from the pial surface to white matter (Hubel and Wiesel, 1959). Studies in cats and monkeys have shown horizontal connections between neurons within a layer (Gilbert, 1983). These horizontal interactions are patchy in nature and connect columns with similar functions. They exist in all cortical layers but are more prominent in superficial (I/II/III) and deep (V/VI) cortical layers (Gilbert, 1983; Rockland and Pandya, 1979). Hubel and Wiesel first observed the structure of ODCs in anesthetized cats during single neuron recording (Hubel and Wiesel, 1962), and later in anesthetized monkeys (Hubel and Wiesel, 1968). In humans, ODCs were first revealed by cytochrome oxidase staining of a post-mortem specimen of a subject who lost an eye prior to death (Horton and Hedley-Whyte, 1984). By examining the distribution of cytochrome oxidase in cortex, they observed a fingerprint-like pattern on the flattened surface with reduced cytochrome oxidase staining in the columns of the missing eye.

Columnar structures are an example of mesoscale neuronal organization that is accessible using ultra-high field MRI systems. ODCs were first imaged in human brain in-vivo using the gradient-echo blood-oxygenation-level-dependent (GRE BOLD) contrast (Ogawa et al., 1990) at 4T with a brief stimulus (Menon et al., 1997). GRE BOLD is highly sensitive to the blood oxygenation change in pre-capillary arterioles, capillaries, venules, and veins (Barth et al., 1999; Frahm et al., 1994; Kim and Ogawa, 2012; Menon, 2012; Menon et al., 1995; Uludağ and Blinder, 2018); however, its poor spatial specificity (Kim et al., 1994; Turner, 2002) hinders the use of this contrast for laminar and columnar studies (Barth and Poser, 2011; Havlicek and Uludağ, 2020; Koopmans et al., 2010; Markuerkiaga et al., 2016; Self et al., 2017). The reasons for this poor spatial specificity are thought to be twofold (Havlicek and Uludağ, 2020). With regard to laminar fMRI, the higher baseline blood volume of the ascending veins in the cortex, and the draining of capillary blood towards the cortical surface through these veins, results in a measured GRE BOLD signal change at each cortical layer that is influenced by the BOLD signal change in that layer plus the signal change from lower cortical layers closer to WM that flows toward the cortical surface. This leads to a degraded laminar signal spatial specificity of this contrast. Correcting for the draining veins effect using a deconvolution approach (Markuerkiaga et al., 2021) or phase regression (Stanley et al., 2021) have been among the post-processing techniques used for laminar BOLD fMRI studies. Additionally, the horizontally extending veins and venules on and below the pial surface contribute to the BOLD signal and result in a horizontal smearing of the columnar signals. Thus, draining venules and veins blur both the laminar and columnar BOLD responses.

Alternatively, imaging contrasts with higher spatial specificity such as Hahn-Spin-Echo (HSE) (Hahn, 1950) BOLD can be employed to study these mesoscale cortical structures (Yacoub et al., 2007). However, the echo-planar readout employed in such studies still contributes to significant large vein contamination as it contains a significant GRE component in addition to the SE term. In recent years, the use of vascular-space-occupancy (VASO) fMRI (Lu et al., 2003) in laminar studies has generated significant interest (Bandettini et al., 2021; Beckett et al., 2020; Chai et al., 2019; Finn et al., 2019; Guidi et al., 2016, 2020; Huber et al., 2014b, 2015, 2016a, 2016b, 2017a, 2021b; Kashyap et al., 2018). VASO is a T_1_ contrast that reflects the extra-vascular tissue signal reduction that occurs as a consequence of the cerebral blood volume (CBV) increase following neuronal activity. As the CBV change is thought to occur mainly in small arterioles and capillaries, the tissue signal reduction around these vessels is expected to be more proximal to the actual site of neural activation (Bollmann and Barth, 2020; Huber et al., 2017b; Lu et al., 2003; Lu and Zijl, 2012). In addition, a simulation study (Akbari et al., 2022) based on a simple cortical vascular model (Markuerkiaga et al., 2016), showed that VASO is less susceptible to large vessel effects compared to BOLD, as blood volume changes in intracortical arteries did not substantially affect the resulting layer-dependent VASO profiles, whereas layer-dependent BOLD profiles showed a bias towards signal contributions from intracortical veins. The feasibility of imaging ODCs in human brain with VASO fMRI has been shown in abstract form (Haenelt et al., 2020). In this work, we compared the VASO and BOLD signal spatial specificities in resolving the ODCs in human V1 at ultra-high magnetic field with mesoscopic (sub-millimeter) isotropic resolution at 7T using a purpose-built V1 radiofrequency (RF) coil. We examined the column-specific VASO and BOLD responses across cortical layers as a function of eye stimulation condition. In addition to resolving the columnar responses during monocular viewing, we examined the columnar responses to binocular viewing for each of the VASO and BOLD contrasts.

## 2 Methods

### 2.1 Participants

Five participants (1 female) with normal or corrected-to-normal vision and no history of neurological impairment were recruited for this study. They gave written informed consent in accordance with and approved by the Human Subjects Research Ethics Board at Western University. In order to identify the dominant eye, all participants were asked to keep both eyes open and focus on a distant object. They then extended one arm out to occlude the object with a finger and closed one eye or the other. The eye that kept the object occluded while the other eye was closed was identified as their dominant eye.

### 2.2 Image acquisition

Imaging was performed at the Center for Functional and Metabolic Mapping at Western University. A head-only 7T scanner (MAGNETOM 7T MRI Plus, Siemens Healthineers AG, Erlangen, Germany) utilizing an AC84 Mark II head-gradient coil with an 8-channel Tx, 32-channel Rx RF coil optimized for occipital-parietal (OP) imaging with no visual obstruction over the face (Gilbert et al., 2017) was used for functional imaging. This OP coil is designed to produce maximum signal-to-noise-ratio (SNR) in the visual cortex, leading to ∼35 % higher temporal signal-to-noise-ratio (tSNR) than a whole-head coil. Anatomic imaging was performed with a whole head RF coil which incorporates an 8-channel transmit dipole coil and 32-channel receive loop coil with an unobstructed visual field (Gilbert et al., 2021). Eight channel parallel transmit was used for all experiments, after B_1_^+^ shimming on the volume of interest.

The phase and magnitude images of the blood-signal-nulled (VASO) and non-nulled (BOLD) contrasts were acquired using the DZNE VASO sequence (Stirnberg and Stöcker, 2021) with a 3D EPI readout (Poser et al., 2010). The phase images were only used for noise reduction using the NORDIC approach (Moeller et al., 2021; Vizioli et al., 2021), as described in the Supplementary materials. NORDIC was not used for the processing of data in the main body of the paper. In each imaging session, five functional runs with 400 volumes each (200 volumes each of the BOLD and VASO contrast) were acquired with the following parameters: *TR/TE/TI_1_/TI_2_* = 3294/21.8/1233/2379 ms, inversion delay = 650 ms, CAIPIRINHA (Breuer et al., 2005) acceleration factor = 3, isotropic voxel size = 0.7 mm, number of slices = 18, partial Fourier in the phase encoding direction= 6/8. The imaging slab was positioned parallel to the calcarine sulcus such that the center of the imaging slab was aligned with the center of the calcarine sulcus, the part of the striate cortex where the ocular dominance columns are located. Whole-brain MP2RAGE (Marques et al., 2010; O’Brien et al., 2014) images with an isotropic resolution of 0.75 mm were acquired for each participant in the same session using the head coil. To minimize the participant’s head motion during the scan, foam cushions were placed around their head.

### 2.3 Experimental set-up and visual stimulus

MR compatible house-made LCD goggles (see Figure 1A) were mounted on the occipital coil above the participant’s eyes that allowed for presenting the stimulus to both eyes (binocular), right eye, and left eye under computer control synchronized to the MRI scanner. To better separate the right and left eye field-of-view, a triangular-shaped septum was placed between the lenses at the nose bridge. Psychopy (Peirce et al., 2019) was used to deliver the stimulus on a screen immediately behind the subject in the scanner using a fibre optic projection system (Avotech SV-6060, Florida, USA). The participants could view the stimulus through a mirror placed over the goggles 28 cm away from the screen, giving an approximately 40-degree field of view.

**Figure 1:**
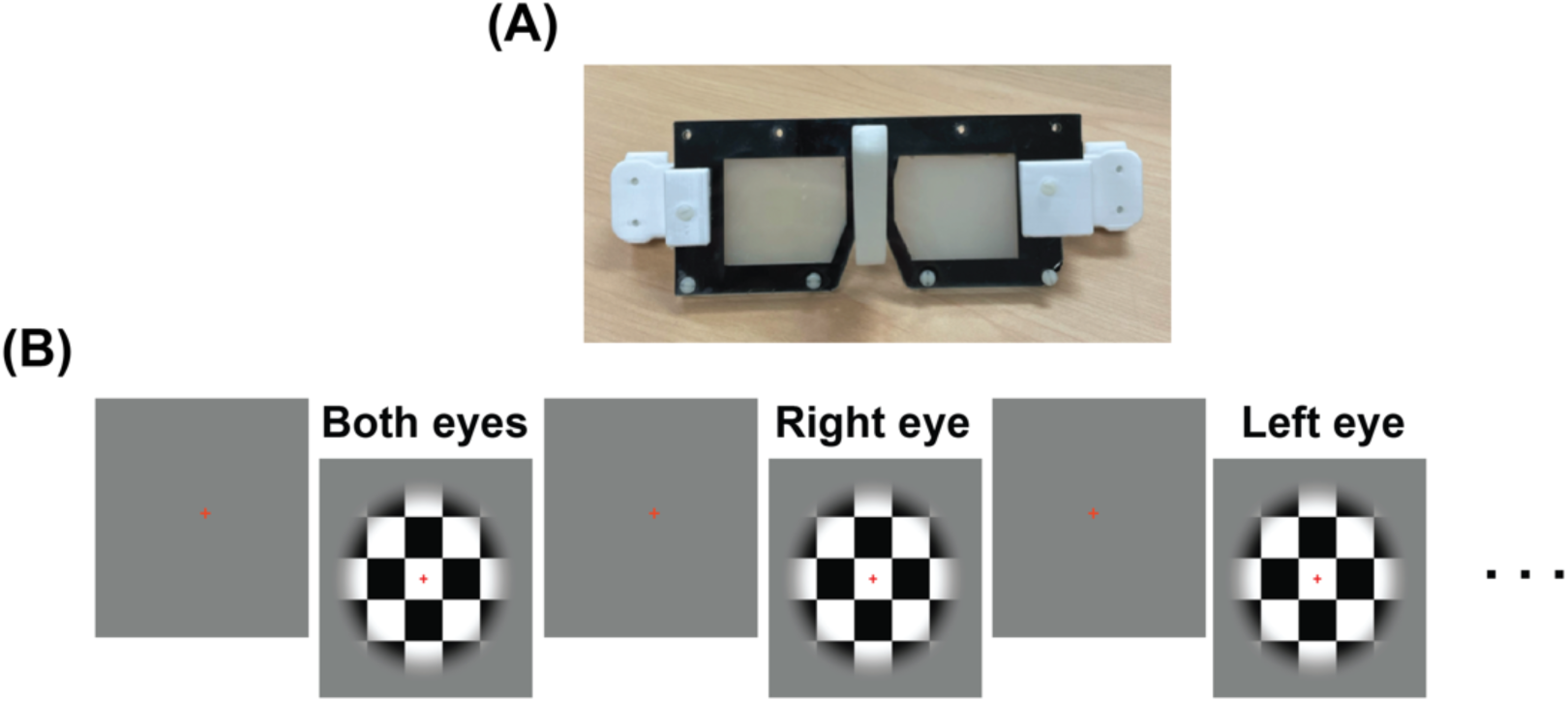
(A) The LCD goggles used to present the stimulus to the participant’s eyes and control the monocular and binocular viewing during the scan. (B) The visual stimulation was consisted of a flickering checkerboard interleaved with a grey screen as the control condition.

The visual stimulus consisted of twenty-two ON- and OFF-blocks of 30 s duration per block. The ON condition was a circular checkerboard (see Figure 1B) contrast-reversing at 10 Hz centered on a grey screen with a red central fixation cross. The checkerboard faded to grey at the edges of the circle to prevent a sharp contrast at the edges. The OFF block was a grey screen with a red central fixation cross.

Each scan started with the grey screen (control condition) presented binocularly, followed by the stimulus presenting binocularly, to the right eye only, and to the left eye only, each interleaved with the control condition. When the stimulus was presented monocularly, the non-stimulated eye viewed an opaque grey lens (see Figure 1A) with the same contrast as the screen presented during the OFF condition. Subjects were instructed to fixate on the red cross at the center of the screen.

We use the term “corresponding eye” to describe the stimulus condition that preferentially activates a column. For example, the corresponding eye for a column determined to have a right eye preference is the right eye and the non-corresponding eye for that column would be the left eye.

### 2.4 Image analysis

#### 2.4.1 GLM analysis

The phase and magnitude images of BOLD and VASO contrasts were first separated based on their DICOM header information using MATLAB (R2022a, The Mathworks, USA) code, and DICOM to NIFTI conversion was performed using dcm2nii (Li et al., 2016). The first volume of each contrast was discarded to ensure *T*_1_ effects were at equilibrium. BOLD and VASO images were motion corrected separately using SPM12 (Wellcome Trust Centre, UK) with a fourth-order B-Spline interpolation. VASO images were BOLD corrected using LAYNII (LN_BOCO) version 2.2.0 (Huber et al., 2021a) to remove the *T^*^_2_* dependency of the extra-vascular signal component. Spatial smoothing was applied along the similar anatomical compartments using LAYNII (LN_GRADSMOOTH) with FWHM = 0.7 mm and selectivity = 0.05. GLM analysis was performed using SPM12 to obtain the BOLD and VASO activation maps for each of binocular, right eye, and left eye stimulus conditions. Voxels with t-values above 3.1 corresponding to an uncorrected significance level of p < 0.001 were identified as the activated regions for both BOLD and BOLD-corrected VASO images. A binary mask created from the region of interest was used as the weighting image in GLM.

#### 2.4.2 Image registration

Presurfer (https://github.com/srikash/presurfer) was used for bias field correction and skull stripping of the MP2RAGE image. The T_1_-EPI image was bias field corrected and denoised using SPM12 and Debian (DenoiseImage) respectively. ITK-SNAP was used to align the MP2RAGE and T_1_-EPI images manually and to obtain an initialization matrix for ANTs registration (Avants et al., 2009). A binary mask was also generated from the region of interest to further improve the registration.

#### 2.4.3 Laminar analysis

The T_1_-EPI image obtained from the BOLD and VASO mean images was used for delineating the GM/CSF and GM/WM boundaries. For smoother layers, the T_1_-EPI image was upsampled to 0.2 mm in-plane resolution using AFNI (3dresample), and then LAYNII was used to create ten equidistant layers across the defined ROI. Note that these layers do not correspond to the histologically defined cortical layers.

To plot the mean signal change across the defined cortical layers, the mean and standard deviation of each of the right eye, left eye, and binocular t-maps were calculated from the layer specific ROIs.

#### 2.4.4 Columnar analysis

To identify the ODCs, voxels from the right eye and left eye t-maps with t-value > 3.1 were identified as the right-eye-dominated and left-eye-dominated columns respectively in BOLD contrast. For VASO, these masks were created with a lower threshold (t > 0) due to the lower sensitivity of this contrast compared to BOLD. These binary masks were then multiplied by the both-eyes t-map to determine left and right eye ODC responses under binocular viewing. These binary masks containing the ODCs were also used to determine the response of a left or right eye column to its corresponding and non-corresponding eye stimulus.

#### 2.4.5 Surface reconstruction

To better visualize the fingerprint pattern of ODCs, a surface flattening analysis pipeline was used. First the MP2RAGE image was registered to the T_1_-EPI image using ANTs (antsApplyTransforms), and then Freesurfer (recon-all) was supplied with the resulting EPI-space anatomical image. Activation maps were registered to the inflated cortical surface using Human Connectome Workbench (Marcus et al., 2011).

## 3 Results

### 3.1 Voxel-wise activation maps and laminar profiles

Figure 2A and 2B show the BOLD and VASO activation maps corresponding to binocular and monocular stimulation overlaid on the reregistered MP2RAGE image for a single representative participant. For VASO, the activated voxels with higher t-values are clearly confined to cortical grey matter, as opposed to BOLD t-maps that show higher activation near the cortical surface. This is particularly noticeable in the sulci. Averaged over all voxels in all subjects, we observed a mean signal change of 3.1 %, 1.7 %, and 1.3 % for binocular, right eye, and left eye stimulation respectively for BOLD. For VASO, the mean signal change across the same stimulus conditions was 0.18 %, 0.08 %, and 0.03 % respectively.

**Figure 2:**
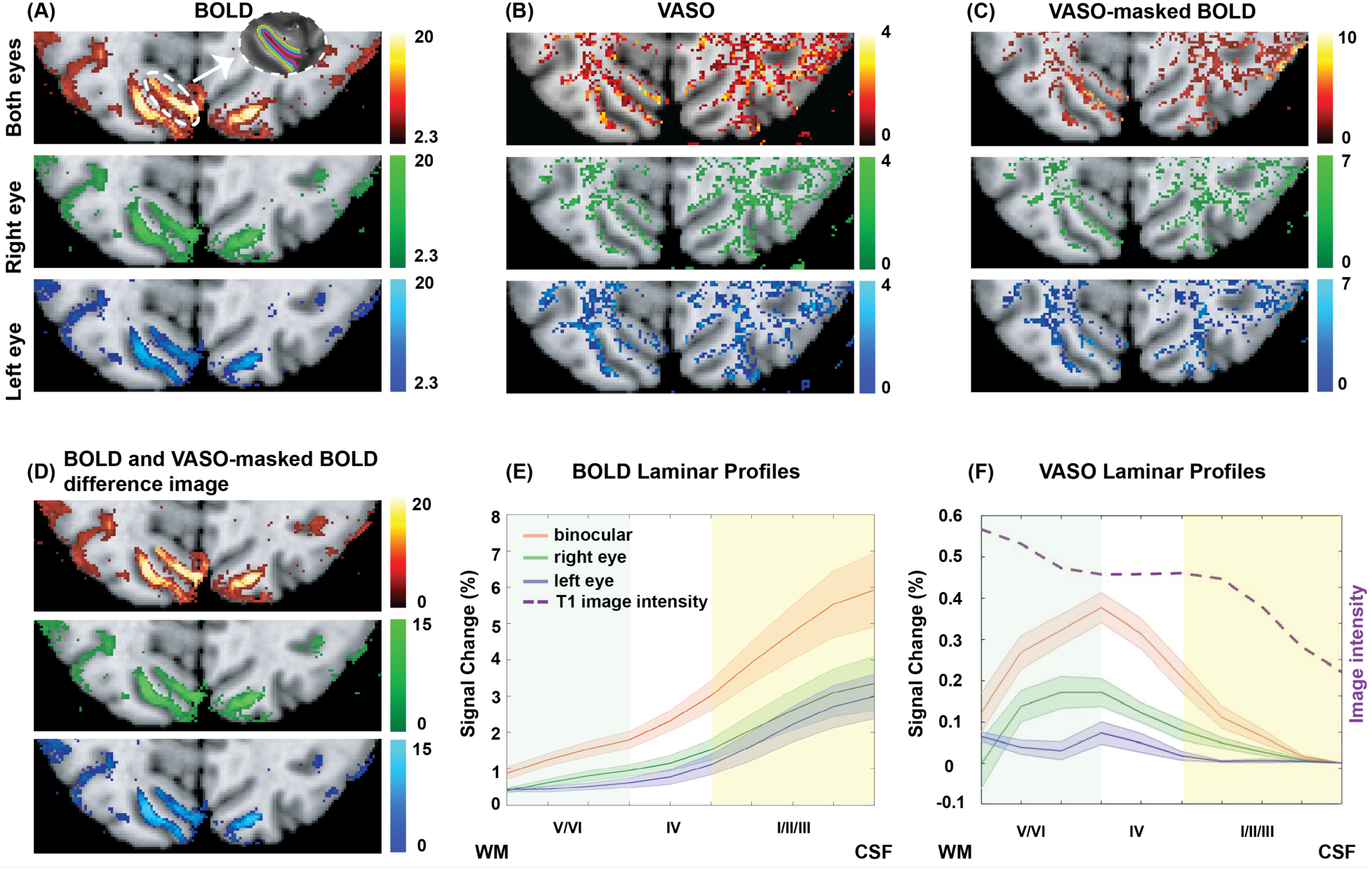
(A) BOLD and (B) VASO activation maps (t-maps) of three stimuli conditions (binocular, right eye, and left eye) overlaid on the MP2RAGE and T_1_-EPI co-registered image for one representative participant. BOLD shows higher activity near cortical surface, while activated voxels in VASO are confined to the grey matter. (C) VASO-masked BOLD activation maps obtained from thresholded VASO t-maps (t > 0) multiplied by the respective BOLD activation maps. An example of the ROI used for our laminar analysis is shown within a dashed white circle in (A). (D) BOLD and VASO-masked BOLD difference image obtained by subtracting the VASO-masked BOLD from BOLD activation maps. (E) BOLD and (F) VASO laminar profiles correspond to all three stimuli averaged across all participants obtained from unthresholded t-maps in (A) and (B). The layer-dependent signal of the T_1_-EPI image is plotted as a reference (dashed purple profile) to identify the approximate location of stria of Gennari. The shaded layer assignments in this figure and all the following figures were approximated from (Henri M Duvernoy et al., 1981).

To facilitate comparison of the BOLD signal from the voxels corresponding only to those identified from the VASO contrast, a VASO-masked BOLD activation map was also created (Figure 2(C)). For these voxels, we observed a mean signal change of 0.27 %, 0.13 %, and 0.14 % for binocular, right eye, and left eye stimulation respectively, comparable to the signal changes seen for the VASO contrast and approximately ten-fold lower than the non-masked BOLD maps. Finally, we subtracted the VASO-masked BOLD voxels from the non-masked BOLD maps, yielding a map of BOLD voxels that are suppressed by using VASO. This map (Figure 2(D)) shows voxels predominantly on the cortical surface where the pial veins are located, which is particularly striking in the sulci.

Using the laminar parcellation approach described in 2.4.3 (see inset in Figure 2(A) as an example), the laminar profiles of BOLD and VASO contrasts corresponding to all three stimulus conditions, averaged across all participants, are shown in Figures 2E and 2F. As expected, the laminar BOLD profiles are skewed towards the location of the large pial veins at the cortical surface, while the laminar VASO profiles peaked in the middle cortical layers, where the thalamic input from LGN arrives in V1.

### 3.2 Ocular dominance column maps

To visualize the ODCs we overlaid the BOLD and VASO activation maps (t-map) on an inflated cortical surface of one representative participant (Figure 3). The result of averaging across all five runs and one run only, are shown in Figure 3A and 3B respectively. The hot (cool) color represents image voxels whose fMRI response was greater in magnitude during right (left) eye stimulation than during left (right) eye stimulation. As shown in Figure 3A, the ∼4.8 × higher sensitivity of the BOLD contrast allows for revealing this map with only one functional run, while VASO contrast required much more averaging to produce a useful map. We note that VASO signal changes are opposite to those for BOLD.

**Figure 3:**
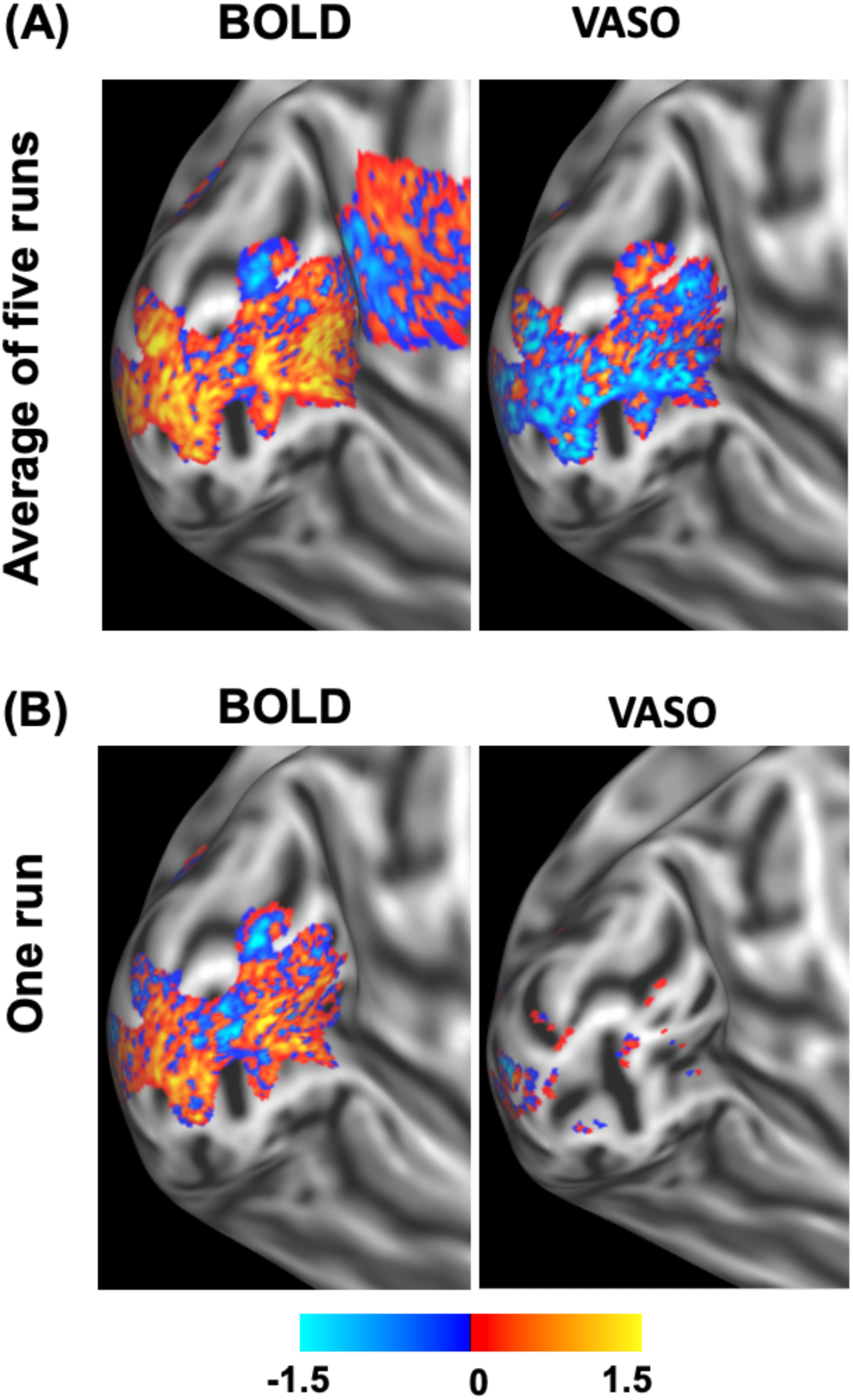
The BOLD and VASO ocular dominance columns maps overlaid on an inflated surface of one representative participant. (A) the activation maps averaged across five runs, and (B) activation map from only one run. The hot (cool) color shows the voxels whose fMRI response was higher in magnitude during the right (left) eye stimulation than the left (right) eye stimulation (i.e., the response of one eye relative to opposite eye).

### 3.3 Right and left eye responses to monocular viewing

Figure 4 demonstrates the laminar dependence of VASO and BOLD responses during monocular viewing. Figure 4(A) depicts the response of the right-eye-dominated-columns to right and left eye stimulation. Conversely, Figure 4(B) shows the response of the left-eye-dominated-columns to the left and right eye stimulation. In all figures, the red (R) and lavender (L) colours are keyed to the right (R) and left (L) eye stimuli respectively. Figures 4(C) and 4(D) show the above-mentioned responses for the VASO contrast.

**Figure 4:**
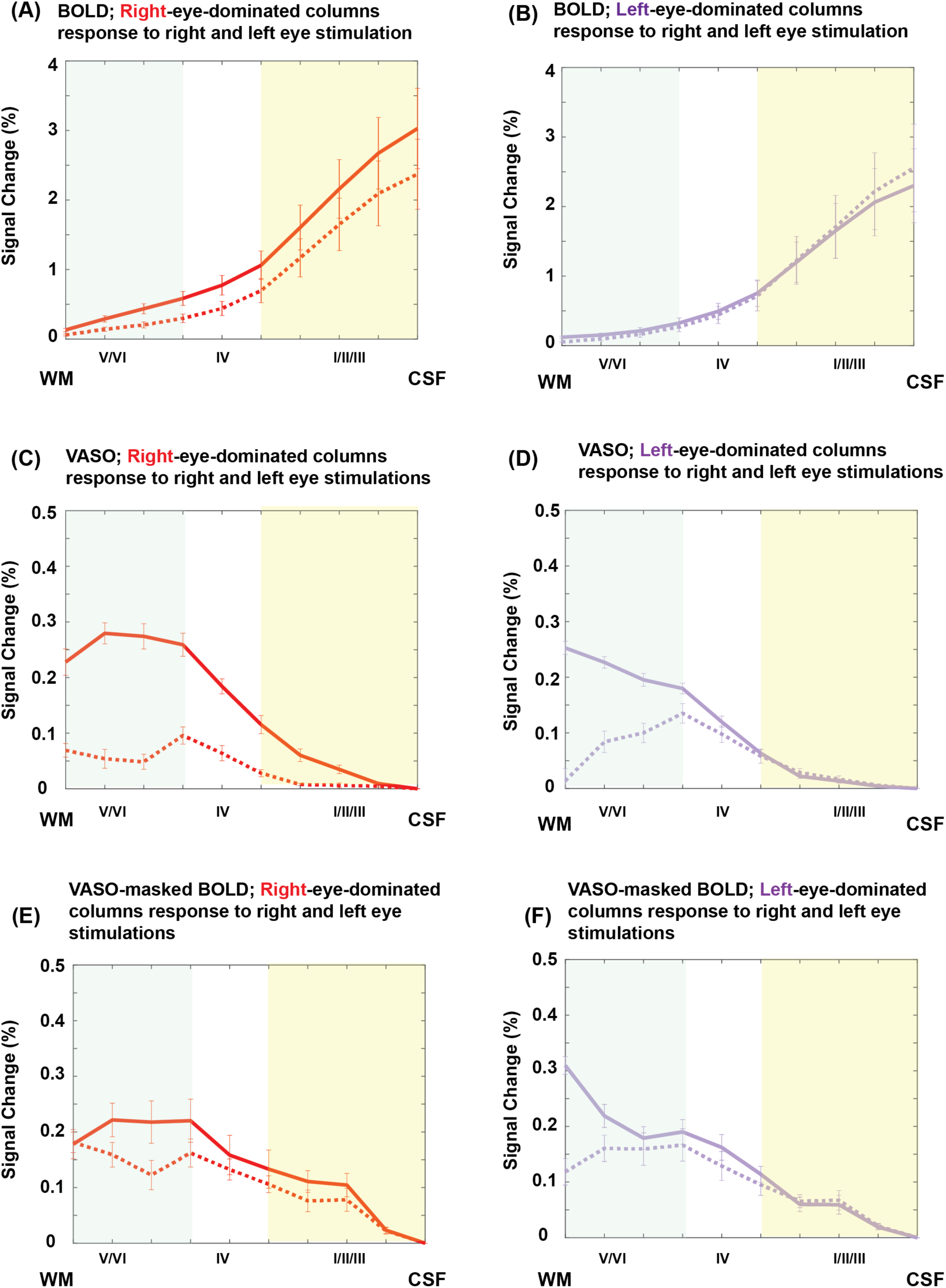
The response of the right-eye-dominated-columns to right (red-solid) and left (red-dashed) eye stimulation in BOLD (A), VASO (C), and (E) VASO-masked BOLD. Similarly, the response of the left-eye-dominated columns to the left (lavender-solid) and right (lavender-dashed) stimulation in BOLD (B), VASO (D), and VASO-masked BOLD (F). Error bars show the standard error of mean. In this figure and the following figures, red (R) and lavender (L) colors refer to right (R) and Left (L) eye responses respectively. The VASO-masked BOLD profiles have been obtained using the columnar masks created from the VASO contrast instead of the BOLD contrast itself.

With VASO, the response of a column to the monocular stimulation of the non- corresponding eye is considerably smaller than that of the corresponding eye, especially for the right eye responses (Figure 4(C)). A paired t-test comparison between the responses of the left (right)-eye-dominated columns to the corresponding and non-corresponding eye stimulation yielded a significant difference for the right-eye-dominated columns (p = 0.003), but not for the left-eye-dominated columns. When examining this difference as a function of layer, significant differences were clear in both columns in the deeper layers. With BOLD, corresponding and non- corresponding eye responses are similar in amplitude and trend across the cortical layers and columns with a greater overlap between the *left* eye (Figure 4(B)) responses.

Figures 4(E) and 4(F) show the right (left)-eye-dominated-columns BOLD laminar responses when using the VASO columnar masks instead of creating the masks from BOLD t-maps themselves (compare with Figures 4(A) and 4(B)). Note the suppressed responses at the cortical surface and the peak of the signal change in deep cortical layers and the similarity in shape to Figure 4(C) and Figure 4(D).

### 3.4 The response of the monocular columns to binocular viewing

Figure 5 shows the response of the right-eye-dominated or left-eye-dominated columns to the corresponding eye stimulation and binocular stimulation. In the deeper layers, the BOLD binocular stimulation response in a column was similar to the monocular response but the BOLD response was considerably higher from the upper-middle to superficial layers (Figure 5(A)). However, for VASO, the binocular stimulation response in a column produced a similar signal change as monocular stimulation (Figure 5(B)). In lower cortical layers, a marginal reduction was visible in the left or right eye columnar responses during monocular stimulation as compared to binocular stimulation. However, a paired t-test didn’t yield a significant difference between these responses in any of the sampled cortical layers. For VASO-masked BOLD (Figure 5(C)), all the responses show a comparable pattern to VASO, however, the response to binocular stimulation was greater than the response to corresponding eye stimulation. Note that the red/lavender solid lines in Figure 5 are the same as the red/lavender solid lines in Figure 4 for the BOLD, VASO, and VASO-masked BOLD analyses.

**Figure 5:**
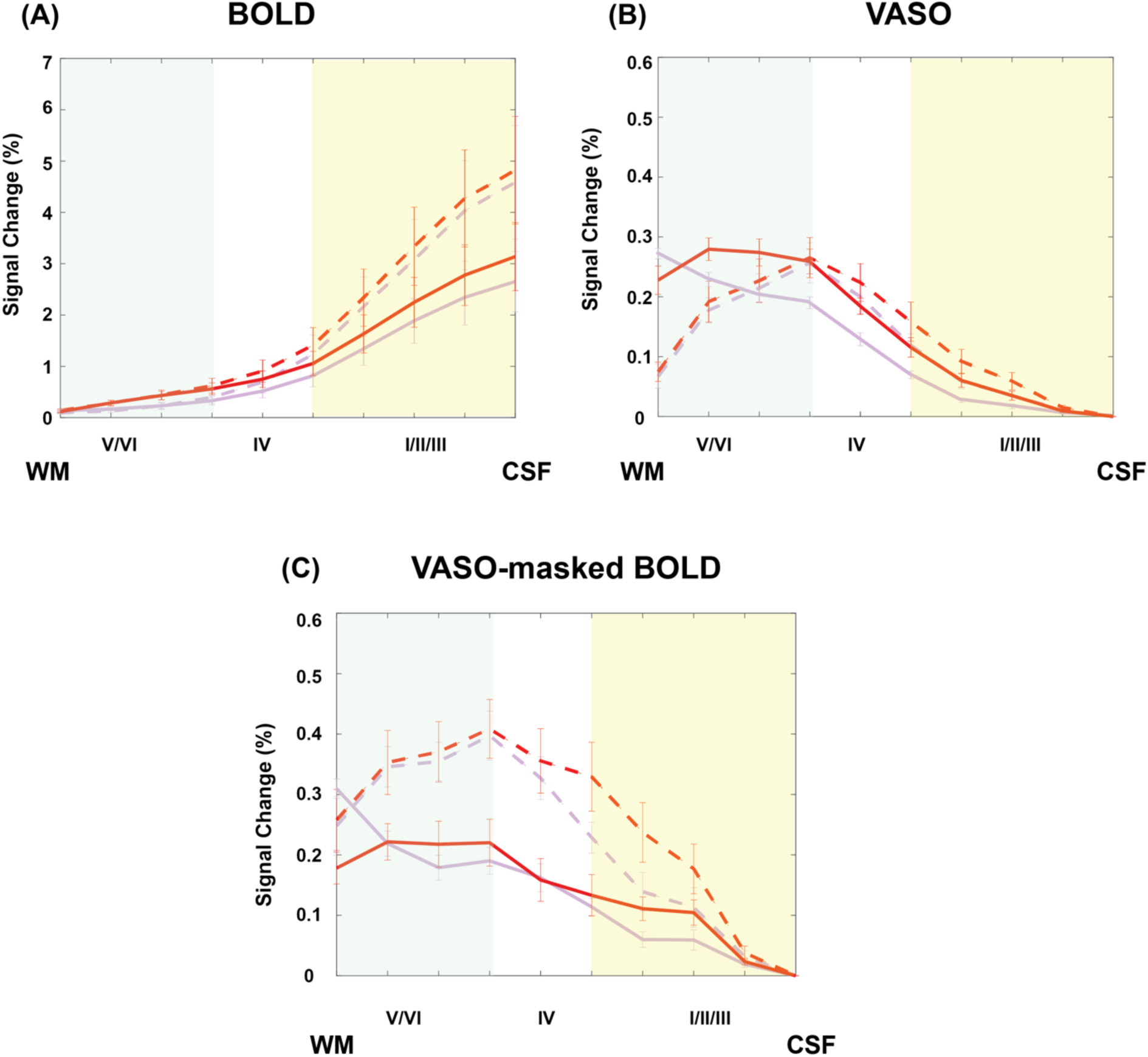
The responses of a left or right eye column to corresponding eye and binocular stimulus presentation for BOLD (A), VASO (B), and VASO-masked BOLD (C). Error bars indicate the standard error of the mean. The solid and dashed red profiles refer to the right-eye-dominated columns response to “right” and “binocular” stimulation respectively. Similarly, the solid and dashed lavender profiles refer to the response of left-eye-dominated columns to “left” and “binocular” stimulation respectively.

## 4 Discussion

### 4.1 Layer vs depth terminology

To be consistent with the prior layer fMRI literature, we have used layer terminology throughout. However, the spatial resolution used in this (0.7 mm isotropic) and prior human studies is completely insufficient to resolve the six histologically defined cortical layers, even when using the standard interpolation schemes. Studies such as this are better considered as cortical depth dependent fMRI. Given that the native EPI resolution can, at best, resolve three layers across the cortex, we have grouped these as superficial, middle, and deep layers (shaded regions in Figure 2(E, F) and assigned the histological layers to these approximately, based on (H.M. Duvernoy et al., 1981).

In this study, we identified ocular dominance columns in healthy human participants at sub-millimeter resolution using BOLD and VASO contrasts at 7T and parcelled the columns into layers. We studied the monocular and binocular responses of the layers in a column under corresponding eye, non-corresponding eye, and binocular stimulation. Instead of using a differential analysis as is common (e.g., left minus right eye response) to isolate columns, we identified columns using a single condition approach, by mapping the response of the corresponding eye to a checkerboard compared to a grey control condition (Dechent and Frahm, 2000). Our results confirm a previous abstract that imaging ocular dominance columns is feasible with VASO; however, due to the lower sensitivity of this contrast, more than one run is required to reveal this structure robustly, making it potentially more sensitive to motion between scans.

The three main findings of this study are: (i) monocular and binocular VASO laminar responses peak in lower middle cortical layers, while BOLD laminar responses were highly skewed towards the cortical surface. (ii) the inter-laminar mixing of the BOLD signal due to the ascending tangential penetrating venules does not significantly blur the layer specific signal characteristics. (iii) Binocular stimulation produces enhanced BOLD responses and decreased VASO responses in a column when compared to corresponding eye stimulation. We discuss each of these findings in detail below.

### 4.2 Monocular and binocular VASO laminar responses peak in lower middle cortical layers, while BOLD laminar responses were highly skewed towards the cortical surface

Figure 2(E) clearly demonstrates the dominance of the pial veins in shaping the BOLD laminar response in all three stimulus conditions. In contrast, and in agreement with previous laminar fMRI studies (Akbari et al., 2022; Huber et al., 2014a; Olman et al., 2012; Polimeni et al., 2010), our layer dependent VASO responses in all three stimulus conditions showed that the VASO contrast was highly effective in suppressing the pial veins that dominate the BOLD contrast (Figure 2(F)). As expected, VASO responses peaked in the middle layers, consistent with the known location of inputs from the thalamus, the known peak of the capillary density (H.M. Duvernoy et al., 1981; Weber et al., 2008a) and the known location of highest neural activity (Kim and Ogawa, 2012; Löwel et al., 1987; Weber et al., 2008b).

We did not see such a peak in the middle cortical layers in the BOLD contrast (Figure 2(E)). This so-called “bump” has been reported in some of the previous studies (Chen et al., 2013; Huber et al., 2017a; Koopmans et al., 2010), however in other studies, including ours, this bump is not visible in the BOLD response profile. This could be due to the insufficient spatial resolution (0.7 mm iso) of the current study and the blurred point spread function introduced by the partial-Fourier 3D-EPI readout that smooths out this local maximum in the laminar BOLD profile (Huber et al., 2018). However, we demonstrate further below that this bump can be recovered in an alternate analysis.

The higher signal spatial specificity of VASO compared to BOLD visible in all laminar profiles reported here likely stem from the intrinsic nature of these fMRI contrasts. BOLD is highly sensitive to the blood oxygenation change in pre-capillary arterioles, capillaries, venules, and venous vasculature and to a lesser degree also depends on the blood volume change in these vessels (Kim and Ogawa, 2012; Uludağ et al., 2009). Since the blood volume change mainly occurs in intracortical arteries, arterioles, capillaries, and to a limited extent in venules (Huber et al., 2014a; Jin and Kim, 2008; Kim et al., 2013, 2007; Kim and Kim, 2010; Zhao et al., 2006), the impact of the CBV change on the BOLD signal is small. This is particularly true on the arterial side, where the susceptibility of the blood matches that of the tissue, and as such, blood volume changes in that part of the vascular tree do not modulate the BOLD signal.

The tangential (to the cortex) ascending intracortical venules drain the deoxygenated blood towards the pial surface, hence, the BOLD signal at upper cortical layer is, in principle, a mix of signals from all lower layers (Havlicek and Uludağ, 2020; Markuerkiaga et al., 2016; Uludağ et al., 2009). This intermixing of the blood draining from each layer potentially reduces the laminar BOLD signal spatial specificity and results in a larger BOLD signal change near the surface, in part because they collect draining blood from the entire depth, and in part because the pial veins comprise a significant fraction of a voxel at these resolutions. Previous studies have attempted to address the draining veins issue by post-processing methods such as masking out the signal from veins (Chen et al., 2013; Koopmans et al., 2010), deconvolution (Hollander et al., 2021; Markuerkiaga et al., 2021), phase regression (Knudsen et al., 2023; Menon, 2002; Stanley et al., 2021) or by employing fMRI contrasts with higher sensitivity to microvasculature such as SE (Goense and Logothetis, 2006; Yacoub et al., 2003; Zhao et al., 2006), GREASE (Beckett et al., 2020; Martino et al., 2013; Polimeni, 2018) and VASO (Huber et al., 2014b; Lu et al., 2003).

We can investigate the role of ascending tangential penetrating venules further using our data, by examining BOLD responses only in voxels contained in the VASO maps (Figure 2(C)). Somewhat surprisingly, the monocular BOLD fractional signal change in the VASO-masked voxels is comparable to the monocular VASO fractional signal change in those voxels, despite the very different signal mechanisms. The VASO- masked BOLD maps clearly exclude the large venous vessels (Figure 2(D)), generating laminar profiles that appear to be spatially concordant between the VASO and VASO- masked BOLD contrasts (compare Figure 4(C, D) to Figure 4(E, F)). The weighting of both these contrasts towards deeper layers is in agreement with previous electrophysiological studies showing that most monocular neurons that preferentially respond to the input from one eye or another are found in middle and deep cortical layers (Dougherty et al., 2019). This result is also consistent with prior high resolution BOLD studies, showing a peak in the middle layers when the pial BOLD signal is masked out (see, for example, Figure 5 in (Chen et al., 2013)). The similarity of the VASO and VASO-masked BOLD laminar profiles in this analysis suggests that inter-laminar mixing of the BOLD signal due to the ascending tangential penetrating venules does not significantly blur the layer specific signal characteristics.

### 4.3 VASO produced better differentiation between stimulated and un-stimulated eye columns compared to BOLD

The BOLD laminar profiles are very similar in amplitude and trend between stimulated and un-stimulated columns and do not show significant separation (Figure 4(A, B)). We speculate that this is due to three characteristics of the draining vein architecture. When monocular neurons are activated upon receiving input from LGN, the tangential ascending intracortical venules draining the deoxygenated blood from those columns can flow through adjacent columns that haven’t received any input, as there is not a one-to-one correspondence between columnar architecture and tangential ascending intracortical venules. In addition, the smaller horizontal (i.e., parallel to the cortical surface) venules feeding into the tangential ascending intracortical venules run horizontally across adjacent columns and will also degrade the spatial specificity. Finally, the pial veins that dominate the GRE BOLD signal form a horizontally diffuse vascular network that does not uniquely correspond to the columnar architecture and hence blurs out the columnar specific activation.

For VASO, the columnar response to the corresponding eye stimulation is greater than their response to non-corresponding eye stimulation (Figure 4(C, D)), at least at lower cortical depths. For VASO, the blood signal is (ideally) nulled, the blood volume changes in the intracortical venules and pial veins are thought to be small, and hence the signal is heavily weighted by the tissue signal around microvasculature within a column. Hence, we expect to see a more column-specific pattern and greater ocular differentiation. The trend of the differentiation between stimulated and un-stimulated columns for VASO, observed in Figure 4(C, D)), is intriguing. The differentiation is poor in the superficial layers but appears to increase in the middle and deep layers. This could be due to the long-range horizontal connections between the columns in the superficial layers, that has been shown previously in cats and monkeys (Berlucchi et al., 1973; Fisken et al., 1975; Gilbert, 1983). These horizontal connections are both inhibitory and faciliatory in nature. They facilitate the connections between the columns with similar functions (e.g., between the right-eye-dominated columns or left-eye-dominated columns) and inhibit the crosstalk between the columns with opposite functions (e.g., between right- and left-eye-dominated columns) (Das and Gilbert, 1995). Thus, neural activity in the superficial layers of a stimulated column can promote activity in the un-stimulated column through these connections, reducing the difference between the columns in these upper layers.

We can examine this behaviour in the VASO-masked BOLD voxels in a similar manner to the laminar analysis. Even though the laminar profiles of VASO and VASO- masked BOLD are quite similar, the segregation of the VASO-masked BOLD response between stimulated and unstimulated columns only marginally improves compared to the pure BOLD activation maps (compare separation of solid versus dashed lines in Figure 4(E, F) and Figure 4(A, B)). While there appears to be a segregation improvement at deep cortical depth (as seen with VASO), these differences are not statistically significant. We speculate that at the spatial resolution of our partial-Fourier 3D-EPI acquisition, within the VASO-masked BOLD columns, there are still horizontal venules that traverse across one or more columns to which BOLD is sensitive even though VASO is not, and these venules could blur the BOLD response between stimulated and un-stimulated columns.

### 4.4 Binocular stimulation produces enhanced BOLD responses and decreased VASO responses in a column when compared to corresponding eye stimulation

Figure 5 presents the BOLD and VASO changes in a column in response to corresponding eye stimulation or binocular stimulation. For the GRE BOLD signal, the superficial layer shows an ∼80% increase in a column when both eyes are stimulated, compared to the corresponding eye monocular stimulation (Figure 5(A)). In the middle and deep layers, no difference between monocular and binocular stimulation is observed. We attribute this increase in the superficial layer during binocular stimulation to the fact that the blood from both active columns pools in the pial veins, causing an increase in the BOLD signal. In the middle and deep layers, the pooling that does occur is in the small horizontal venules that run across columns. While one might expect some degree of enhanced response, the VASO signal suggests that there is inhibition of the activity in a column in the deep layer during binocular stimulation, and this may account for the lack of additive enhancement.

For VASO, the columnar response to binocular stimulation was comparable to monocular stimulation of the corresponding eye in the superficial and middle layers (Figure 5(B)). This indicates that the blood volume changes that VASO is sensitive to are well confined within the columns with different eye-specific inputs at both the pial surface and the intracortical vessels in the superficial and middle layers. When both eyes are receiving inputs, the VASO signal appears to remain specific to the input from stimulated columns in these layers. As the corresponding/non-corresponding eye stimulation results show, there are no detectable blood volume changes in the horizontal venules that propagate between an active and an inactive column. In the lower middle and deep layers, the binocular stimulus condition produces a reduced VASO response compared to monocular stimulation of the corresponding eye. We speculate that this is due to inhibitory interneurons in these layers.

For VASO-masked BOLD, even though the pattern and amplitude of columnar responses to binocular and corresponding eye monocular stimulation were comparable to VASO, the columnar response to binocular viewing was significantly greater than the corresponding eye response (Figure 5(C)). The VASO columnar mask improved the BOLD signal spatial specificity by suppressing pial vein effects; however, as mentioned earlier, the BOLD signal leakage between adjacent columns through small venules running parallel to surface, still led to a greater response when the stimulus was presented binocularly.

### 4.5 Comparison with earlier studies

The majority of ODC imaging with fMRI have been performed using GRE BOLD contrast, though its intrinsically limited signal spatial specificity makes the interpretation of depth-specific analysis challenging. Early ODC fMRI maps (Dechent and Frahm, 2000; Goodyear et al., 2009; Goodyear and Menon, 2001) were acquired with sub-millimetre in-plane resolution and a few millimetres resolution in slice direction, which is not suitable for cortical depth analysis. Later on, the wider availability of ultra-high magnetic field scanners (≥ 7T) and advances in hardware developments, enabled studying the brain structure’s function with sub-millimeter isotropic resolution. This was promising for investigating the layer dependent, column-specific cortical processing such as the feedforward and feedback responses. The higher magnetic field also allowed employing imaging contrasts with higher signal spatial specificity compared to BOLD, but at the cost of lower sensitivity. Yacob et al., acquired the human ODCs fMRI maps using SE and GRE contrast at 7T (Yacoub et al., 2007), showing the superior SE signal specificity for resolving these fine-scale human brain structures. Zaretskaya et al., examined the eye-selective responses of the GRE BOLD signal at 3T and 9.4T (Zaretskaya et al., 2020) and showed that the higher sensitivity and resolution at 9.4T allowed investigation of the eye-selective responses in a depth-dependent manner, however, the eye-selectivity was maximum at cortical surface, as the laminar GRE BOLD response is biased towards the location of the large veins. Hollander et al., employed a deconvolution approach to dissociate the thalamic feedforward and feedback responses in human V1 (Hollander et al., 2021), and they showed when correcting for the draining vein effect, the laminar BOLD response peaks at deep and middle cortical layers, as we observe in the VASO-masked BOLD voxels.

### 4.6 Limitations and outlook

One limitation of this study is the relatively low in-plane spatial resolution of the 3D EPI sequence. The 0.7 mm resolution is further blurred by a point spread function arising from the T_2_^*^ decay during the EPI readout. This spatial resolution is similar to the size of human ocular dominance columns (Adams et al., 2007; Horton and Hedley-Whyte, 1984; Menon and Goodyear, 1999). The dominant eye (usually the right eye) would suffer slightly less from partial volume effects due to inter-columnar draining veins because the dominant eye columns are slightly larger (∼1 mm) which may explain why right eye dominant columns display a greater response differential to left eye stimulation (Figure 4(A)) while the smaller left eye columns (∼0.7 mm) show little difference between left and right eye stimulation (Figure 4(B)). This draining vein contamination between columns is mostly eliminated in VASO (Figure 4C/D).

The current work, for the first time shows the depth-specific monocular and binocular responses of human ODCs using VASO fMRI at ultra-high magnetic field. The isotropic sub-millimeter resolution allowed us to explore the fMRI signal change across cortical depth, and the higher signal spatial specificity of the VASO contrast helped to segregate the columnar responses. Future work may benefit from the high sensitivity of the ultra-high magnetic field in addition to the higher signal specificity of the VASO contrast to map the spatial feature and orientation processing of the orientation columns in addition to the ocular dominance columns in human V1.

Caution needs to be exercised in the interpretation of signal changes in BOLD at the mesoscale both in terms of feedforward/feedback effects and inhibitory and excitatory effects. The interpretation of the results at the submillimetre resolution heavily depends on the level of signal spatial specificity and sensitivity of the fMRI contrast used. For example, Figure 4 demonstrates that the BOLD signal in a column does not discriminate well between stimulation of the corresponding eye and the non-corresponding eye. VASO on the other hand indicates that there is a good separation between these two stimulus conditions in a column in the middle and deep layers. The interpretation of what the underlying neurons are doing is very different in these two contrasts. Similarly, the BOLD signal in Figure 5 would suggest that there is an additive effect in the superficial layer when both eyes are stimulated, compared to one eye. However, the VASO data shown no such additive effect in the superficial layer, and indeed shows a decrease during binocular stimulation in the middle and deep layers. Again, the interpretation of underlying neural activity would be very different in these two contrasts. In both these examples, VASO is more consistent with what is known about the laminar inputs to ODCs and the nature of the horizontal connections between the ODCs.

However, the SNR for BOLD is significantly higher. VASO is a noisy contrast and needs an extensive amount of averaging which leads to a longer scan time, and consequently subject discomfort and motion. This makes the use of this contrast challenging in patient populations. Recently, a thermal noise reduction scheme (NORDIC) has been introduced (Moeller et al., 2021; Vizioli et al., 2021) that could be useful to apply in noisy fMRI contrasts like VASO. In the supplementary materials of this manuscript, we have provided the depth-dependent, column-specific results presented in this manuscript after applying NORDIC.

## 5 Conclusion

In summary, we studied the depth dependent BOLD and VASO monocular and binocular responses in human ocular dominance columns at 7T with sub-millimetre isotropic resolution. Our results show that ocular segregation (fMRI signal differentiability between the adjacent columns) is better achieved with VASO, as the blood volume changes are confined to the microvasculature both within and across the depth of a column. On the other hand, the GRE BOLD signal extends across columns and layers due to the distributed, web-like nature of the draining venules and veins. It is further confounded by the pial veins, which dominate the BOLD signal response and also mix the signals from multiple columns.

These results also suggest that despite very different underlying biophysical contrast mechanisms, GRE BOLD *could* have a similar spatial specificity to highly localized neuronal activity as VASO, provided that *both* the horizontal and tangential venous vessel GRE BOLD contributions in the cortex can be suppressed (as VASO does).

## 6 Acknowledgement

The authors would like to thank Trevor Szekeres and Scott Charlton for their assistance with data acquisition. We also acknowledge the helpful discussions with Peter Bandettini and Renzo Huber. This work was supported by a CIHR Foundation grant, BrainsCAN-Canada First Research Excellence Fund, and a Brain Canada Platform Support Grant.

## 8 Data Availability Statement

The raw data are available upon request.

## References

Adams, D.L., Sincich, L.C., Horton, J.C., 2007. Complete Pattern of Ocular Dominance Columns in Human Primary Visual Cortex. J Neurosci 27, 10391–10403. https://doi.org/10.1523/jneurosci.2923-07.2007

Akbari, A., Bollmann, S., Ali, T.S., Barth, M., 2022. Modelling the depth-dependent VASO and BOLD responses in human primary visual cortex. Hum. Brain Mapp. https://doi.org/10.1002/hbm.26094

Avants, B., Tustison, N.J., Song, G., 2009. Advanced Normalization Tools: V1.0. Insight J. https://doi.org/10.54294/uvnhin

Bandettini, P.A., Huber, L., Finn, E.S., 2021. Challenges and opportunities of mesoscopic brain mapping with fMRI. Current Opinion in Behavioral Sciences 40, 189–200. https://doi.org/10.1016/j.cobeha.2021.06.002

Barth, M., Poser, B.A., 2011. Advances in high-field BOLD fMRI. Mateials. 4, 1941–1955. https://doi.org/10.3390/ma4111941

Barth, M., Reichenbach, J.R., Venkatesan, R., Moser, E., Haacke, E.M., 1999. High-resolution, multiple gradient-echo functional MRI at 1.5 T. Magn Reson Imaging 17, 321–329. https://doi.org/10.1016/s0730-725x(98)00191-x

Beckett, A.J., Dadakova, T., Townsend, J., Huber, L., Park, S., Feinberg, D.A., 2020. Comparison of BOLD and CBV using 3D EPI and 3D GRASE for cortical layer functional MRI at 7 T. Magnetic Mesonance in Medicine. 84, 3128–3145. https://doi.org/10.1002/mrm.28347

Berlucchi, G., Brindley, G.S., Brooks, B., Creutzfeldt, O.D., Dodt, E., Doty, R.W., Freund, H.-J., Gross, C.G., Jeffreys, D.A., Jung, R., Kuhnt, U., MacKay, D.M., Marg, E., Negrão, N., Rizzolatti, G., Sprague, J.M., Székely, G., Szentágothai, J, Whitteridge, D., Szentágothai, János, 1973. Visual Centers in the Brain. Handb Sens Physiology 269–324. https://doi.org/10.1007/978-3-642-65495-4_8

Bollmann, S., Barth, M., 2020. New acquisition techniques and their prospects for the achievable resolution of fMRI. Prog Neurobiol 207, 101936. https://doi.org/10.1016/j.pneurobio.2020.101936

Breuer, F.A., Blaimer, M., Heidemann, R.M., Mueller, M.F., Griswold, M.A., Jakob, P.M., 2005. Controlled aliasing in parallel imaging results in higher acceleration (CAIPIRINHA) for multi-slice imaging. Magnet Reson Med 53, 684–691. https://doi.org/10.1002/mrm.20401

Chai, Y., Li, L., Huber, L., Poser, B.A., Bandettini, P.A., 2019. Integrated VASO and perfusion contrast: A new tool for laminar functional MRI. Neuroimage. 116358. https://doi.org/10.1016/j.neuroimage.2019.116358

Chen, G., Wang, F., Gore, J.C., Roe, A.W., 2013. Layer-specific BOLD activation in awake monkey V1 revealed by ultra-high spatial resolution functional magnetic resonance imaging. Neuroimage. 64, 147–155. https://doi.org/10.1016/j.neuroimage.2012.08.060

Das, A., Gilbert, C.D., 1995. Long-range horizontal connections and their role in cortical reorganization revealed by optical recording of cat primary visual cortex. Nature 375, 780– 784. https://doi.org/10.1038/375780a0

Dechent, P., Frahm, J., 2000. Direct mapping of ocular dominance columns in human primary visual cortex. Neuroreport 11, 3247–3249. https://doi.org/10.1097/00001756-200009280-00039

Dougherty, K., Cox, M.A., Westerberg, J.A., Maier, A., 2019. Binocular Modulation of Monocular V1 Neurons. Curr Biol 29, 381–391.e4. https://doi.org/10.1016/j.cub.2018.12.004

Douglas, R.J., Martin, K.A., 2004. Neuronal circuits of the neocortex. Annu. Rev. Neurosci. 27, 419–451. https://doi.org/10.1146/annurev.neuro.27.070203.144152

Duvernoy, Henri M, Delon, S., Vannson, J., 1981. Cortical blood vessels of the human brain. Brain Research Bulletin. 7, 519–579. https://doi.org/10.1016/0361-9230(81)90007-1

Duvernoy, H.M., Delon, S., Vannson, J.L., 1981. Cortical blood vessels of the human brain. Brain Res Bull 7, 519–579. https://doi.org/10.1016/0361-9230(81)90007-1

Felleman, D.J., Van, D.E., 1991. Distributed hierarchical processing in the primate cerebral cortex. Cerebral Cortex. 1, 1–47. https://doi.org/10.1093/cercor/1.1.1-a

Finn, E.S., Huber, L., Jangraw, D.C., Molfese, P.J., Bandettini, P.A., 2019. Layer-dependent activity in human prefrontal cortex during working memory. Nature Neuroscience. 22, 1687–1695. https://doi.org/10.1038/s41593-019-0487-z

Fisken, R.A., Garey, L.J., Powell, T.P.S., 1975. The intrinsic, association and commissural connections of area 17 of the visual cortex. Philosophical Transactions Royal Soc Lond B Biological Sci 272, 487–536. https://doi.org/10.1098/rstb.1975.0099

Frahm, J., Merboldt, K., Hänicke, W., Kleinschmidt, A., Boecker, H., 1994. Brain or veinIoxygenation or flow? On signal physiology in functional MRI of human brain activation. Nmr Biomed. 7, 45–53. https://doi.org/10.1002/nbm.1940070108

Gilbert, C.D., 1983. Microcircuitry of the Visual Cortex. Annu Rev Neurosci 6, 217–247. https://doi.org/10.1146/annurev.ne.06.030183.001245

Gilbert, K.M., Gati, J.S., Menon, R.S., 2017. Occipital-Parietal Coil with variable-density element distribution for 7T functional imaging. Presented at the Proceedings of the 25th International Society for Magnetic Resonance in Medicine Annual Meeting. Honolulu, USA. https://doi.org/https://cds.ismrm.org/protected/17MProceedings/PDFfiles/4307.html

Gilbert, K.M., Klassen, L.M., Mashkovtsev, A., Zeman, P., Menon, R.S., Gati, J.S., 2021. Radiofrequency coil for routine ultra-high-field imaging with an unobstructed visual field. Nmr Biomed 34, e4457. https://doi.org/10.1002/nbm.4457

Goense, J.B., Logothetis, N.K., 2006. Laminar specificity in monkey V1 using high-resolution SE-fMRI. Magnetic Resonance Imaging. 24, 381–392. https://doi.org/10.1016/j.mri.2005.12.032

Goodyear, B.G., Menon, R.S., 2001. Brief visual stimulation allows mapping of ocular dominance in visual cortex using fMRI. Hum Brain Mapp 14, 210–217. https://doi.org/10.1002/hbm.1053

Goodyear, B.G., Nicolle, D.A., Menon, R.S., 2009. High resolution fMRI of ocular dominance columns within the visual cortex of human amblyopes. Strabismus 10, 129–136. https://doi.org/10.1076/stra.10.2.129.8140

Guidi, M., Huber, L., Lampe, L., Gauthier, C.J., Möller, H.E., 2016. Lamina-dependent calibrated BOLD response in human primary motor cortex. Neuroimage 141, 250–261. https://doi.org/10.1016/j.neuroimage.2016.06.030

Guidi, M., Huber, L., Lampe, L., Merola, A., Ihle, K., Möller, H.E., 2020. Cortical laminar resting-state signal fluctuations scale with the hypercapnic blood oxygenation level-dependent response. Hum Brain Mapp 41, 2014–2027. https://doi.org/10.1002/hbm.24926

Haenelt, D., Weiskopf, N., Vaculciakova, L., Mueller, R., Nasr, S., Polimeni, J., Tootell, R., Huber, L., Sereno, M., Trampel, R., 2020. Mapping ocular dominance columns in humans using GE-EPI, SE-EPI and SS-SI-VASO at 7 T, in: The International Society for Magnetic Resonance in Medicine. https://doi.org/https://submissions2.mirasmart.com/ISMRM2020/ViewSubmission.aspx?sbmID=2785&;validate=false

Hahn, E.L., 1950. Spin Echoes. Phys Rev 80, 580–594. https://doi.org/10.1103/physrev.80.580

Havlicek, M., Uludağ, K., 2020. A dynamical model of the laminar BOLD response. NeuroImage. 204, 116209. https://doi.org/10.1016/j.neuroimage.2019.116209

Henderickson, A.E., Wilson, J.R., Ogren, M.P., 1978. The neurological organization of pathways between the dorsal lateral geniculate nucleus and visual cortex in old world and new world primates. J Comp Neurol 182, 123–136. https://doi.org/10.1002/cne.901820108

Hollander, G. de, Zwaag, W. van der, Qian, C., Zhang, P., Knapen, T., 2021. Ultra-high field fMRI reveals origins of feedforward and feedback activity within laminae of human ocular dominance columns. Neuroimage 228, 117683. https://doi.org/10.1016/j.neuroimage.2020.117683

Horton, J.C., Hedley-Whyte, E.T., 1984. Mapping of cytochrome oxidase patches and ocular dominance columns in human visual cortex. Philosophical Transactions Royal Soc Lond B Biological Sci 304, 255–272. https://doi.org/10.1098/rstb.1984.0022

Hubel, D.H., Wiesel, T.N., 1968. Receptive fields and functional architecture of monkey striate cortex. J Physiology 195, 215–243. https://doi.org/10.1113/jphysiol.1968.sp008455

Hubel, D.H., Wiesel, T.N., 1962. Receptive fields, binocular interaction and functional architecture in the cat’s visual cortex. J Physiology 160, 106–154. https://doi.org/10.1113/jphysiol.1962.sp006837

Hubel, D.H., Wiesel, T.N., 1959. Receptive fields of single neurones in the cat’s striate cortex. J Physiology 148, 574–591. https://doi.org/10.1113/jphysiol.1959.sp006308

Huber, L., Goense, J., Kennerley, A.J., Ivanov, D., Krieger, S.N., Lepsien, J., Trampel, R., Turner, R., Möller, H.E., 2014a. Investigation of the neurovascular coupling in positive and negative BOLD responses in human brain at 7 T. Neuroimage 97, 349–362. https://doi.org/10.1016/j.neuroimage.2014.04.022

Huber, L., Goense, J., Kennerley, A.J., Trampel, R., Guidi, M., Reimer, E., Ivanov, D., Neef, N., Gauthier, C.J., Turner, R., 2015. Cortical lamina-dependent blood volume changes in human brain at 7 T. Neuroimage. 107, 23–33. https://doi.org/10.1016/j.neuroimage.2014.11.046

Huber, L., Handwerker, D.A., Jangraw, D.C., Chen, G., Hall, A., Stüber, C., Gonzalez-Castillo, J., Ivanov, D., Marrett, S., Guidi, M., 2017a. High-resolution CBV-fMRI allows mapping of laminar activity and connectivity of cortical input and output in human M1. Neuron. 96, 1253–1263. e7. https://doi.org/10.1016/j.neuron.2017.11.005

Huber, L., Ivanov, D., Guidi, M., Turner, R., Uludağ, K., Möller, H.E., Poser, B.A., 2016a. Functional cerebral blood volume mapping with simultaneous multi-slice acquisition. Neuroimage 125, 1159–1168. https://doi.org/10.1016/j.neuroimage.2015.10.082

Huber, L., Ivanov, D., Handwerker, D.A., Marrett, S., Guidi, M., Uludağ, K., Bandettini, P.A., Poser, B.A., 2018. Techniques for blood volume fMRI with VASO: From low-resolution mapping towards sub-millimeter layer-dependent applications. Neuroimage 164, 131–143. https://doi.org/https://doi.org/10.1016/j.neuroimage.2016.11.039

Huber, L., Ivanov, D., Handwerker, D.A., Marrett, S., Guidi, M., Uludağ, K., Bandettini, P.A., Poser, B.A., 2016b. Techniques for blood volume fMRI with VASO: from low-resolution mapping towards sub-millimeter layer-dependent applications. Neuroimage 164, 131–143. https://doi.org/10.1016/j.neuroimage.2016.11.039

Huber, L., Ivanov, D., Krieger, S.N., Streicher, M.N., Mildner, T., Poser, B.A., Möller, H.E., Turner, R., 2014b. Slab-selective, BOLD-corrected VASO at 7 Tesla provides measures of cerebral blood volume reactivity with high signal-to-noise ratio. Magnetic Resonance in Medicine. 72, 137–148. https://doi.org/10.1002/mrm.24916

Huber, L., Uludağ, K., Möller, H.E., 2017b. Non-BOLD contrast for laminar fMRI in humans: CBF, CBV, and CMRO2. Neuroimage. https://doi.org/10.1016/j.neuroimage.2017.07.041

Huber, L.R., Poser, B.A., Bandettini, P.A., Arora, K., Wagstyl, K., Cho, S., Goense, J., Nothnagel, N., Morgan, A.T., Hurk, J. van den, 2021a. LAYNII: a software suite for layer-fMRI. Neuroimage 237, 118091. https://doi.org/10.1016/j.neuroimage.2021.118091

Huber, L.R., Poser, B.A., Kaas, A.L., Fear, E.J., Dresbach, S., Berwick, J., Goebel, R., Turner, R., Kennerley, A.J., 2021b. Validating layer-specific VASO across species. NeuroImage 118195. https://doi.org/https://doi.org/10.1016/j.neuroimage.2021.118195

Jin, T., Kim, S.-G., 2008. Cortical layer-dependent dynamic blood oxygenation, cerebral blood flow and cerebral blood volume responses during visual stimulation. Neuroimage. 43, 1–9. https://doi.org/10.1016/j.neuroimage.2008.06.029

Kashyap, S., Ivanov, D., Havlicek, M., Sengupta, S., Poser, B.A., Uludağ, K. %J S. reports, 2018. Resolving laminar activation in human V1 using ultra-high spatial resolution fMRI at 7T. Scientific Reports. 8, 1–11. https://doi.org/https://doi.org/10.1038/s41598-018-35333-3

Kim, S., Harel, N., Jin, T., Kim, T., Lee, P., Zhao, F., 2013. Cerebral blood volume MRI with intravascular superparamagnetic iron oxide nanoparticles. NMR in Biomedicine. 26, 949– 962. https://doi.org/10.1002/nbm.2885

Kim, S., Hendrich, K., Hu, X., Merkle, H., Uǧurbil, K., 1994. Potential pitfalls of functional MRI using conventional gradient-recalled echo techniques. NMR in Biomedicine. 7, 69–74. https://doi.org/10.1002/nbm.1940070111

Kim, S.-G., Ogawa, S., 2012. Biophysical and Physiological Origins of Blood Oxygenation Level-Dependent fMRI Signals. J Cereb Blood Flow Metabolism 32, 1188–1206. https://doi.org/10.1038/jcbfm.2012.23

Kim, T., Hendrich, K.S., Masamoto, K., Kim, S.-G., 2007. Arterial versus total blood volume changes during neural activity-induced cerebral blood flow change: implication for BOLD fMRI. Journal of Cerebral Blood Flow & Metabolism 27, 1235–1247. https://doi.org/10.1038/sj.jcbfm.9600429

Kim, T., Kim, S.-G., 2010. Cortical layer-dependent arterial blood volume changes: improved spatial specificity relative to BOLD fMRI. Neuroimage. 49, 1340–1349. https://doi.org/10.1016/j.neuroimage.2009.09.061

Knudsen, L., Bailey, C.J., Blicher, J.U., Yang, Y., Zhang, P., Lund, T.E., 2023. Improved sensitivity and microvascular weighting of 3T laminar fMRI with GE-BOLD using NORDIC and phase regression. Neuroimage 271, 120011. https://doi.org/10.1016/j.neuroimage.2023.120011

Koopmans, P.J., Barth, M., Norris, D.G., 2010. Layer-specific BOLD activation in human V1. Human Brain Mapping. 31, 1297–1304. https://doi.org/10.1002/hbm.20936

Li, X., Morgan, P.S., Ashburner, J., Smith, J., Rorden, C., 2016. The first step for neuroimaging data analysis: DICOM to NIfTI conversion. J Neurosci Meth 264, 47–56. https://doi.org/10.1016/j.jneumeth.2016.03.001

Löwel, S., Freeman, B., Singer, W., 1987. Topographic organization of the orientation column system in large flat-mounts of the cat visual cortex: A 2-deoxyglucose study. J Comp Neurol 255, 401–415. https://doi.org/10.1002/cne.902550307

Lu, H., Golay, X., Pekar, J.J., Zijl, P. van, 2003. Functional magnetic resonance imaging based on changes in vascular space occupancy. Magnetic Resonance in Medicine. 50, 263–274. https://doi.org/10.1002/mrm.10519

Lu, H., Zijl, P.C. van, 2012. A review of the development of Vascular-Space-Occupancy (VASO) fMRI. Neuroimage. 62, 736–742. https://doi.org/10.1016/j.neuroimage.2012.01.013

Marcus, D.S., Harwell, J., Olsen, T., Hodge, M., Glasser, M.F., Prior, F., Jenkinson, M., Laumann, T., Curtiss, S.W., Essen, D.C.V., 2011. Informatics and Data Mining Tools and Strategies for the Human Connectome Project. Front Neuroinform 5, 4. https://doi.org/10.3389/fninf.2011.00004

Markuerkiaga, I., Barth, M., Norris, D.G., 2016. A cortical vascular model for examining the specificity of the laminar BOLD signal. Neuroimage. 132, 491–498. https://doi.org/10.1016/j.neuroimage.2016.02.073

Markuerkiaga, I., Marques, J.P., Gallagher, T.E., Norris, D.G., 2021. Estimation of laminar BOLD activation profiles using deconvolution with a physiological point spread function. Journal of Neuroscience Methods. 353, 109095. https://doi.org/10.1016/j.jneumeth.2021.109095

Marques, J.P., Kober, T., Krueger, G., Zwaag, W. van der, Moortele, P.-F.V. de, Gruetter, R., 2010. MP2RAGE, a self bias-field corrected sequence for improved segmentation and T1- mapping at high field. Neuroimage. 49, 1271–1281. https://doi.org/10.1016/j.neuroimage.2009.10.002

Martino, F.D., Zimmermann, J., Muckli, L., Ugurbil, K., Yacoub, E., Goebel, R., 2013. Cortical depth dependent functional responses in humans at 7T: improved specificity with 3D GRASE. PLoS One 8, e60514. https://doi.org/10.1371/journal.pone.0060514

Menon, R.S., 2012. The great brain versus vein debate. Neuroimage 62, 970–974. https://doi.org/10.1016/j.neuroimage.2011.09.005

Menon, R.S., 2002. Postacquisition suppression of large-vessel BOLD signals in high-resolution fMRI. Magn. Reson. Med. 47, 1–9. https://doi.org/10.1002/mrm.10041

Menon, R.S., Goodyear, B.G., 1999. Submillimeter functional localization in human striate cortex using BOLD contrast at 4 Tesla: Implications for the vascular point-spread function. Magn. Reson. Med. 41, 230–235. https://doi.org/10.1002/(sici)1522-2594(199902)41:2≤230::aid-mrm3>3.0.co;2-o

Menon, R.S., Ogawa, S., Hu, X., Strupp, J.P., Anderson, P., Uǧurbil, K., 1995. BOLD based functional MRI at 4 Tesla includes a capillary bed contribution: echo-planar imaging correlates with previous optical imaging using intrinsic signals. Magnetic Resonance in Medicine. 33, 453–459. https://doi.org/10.1002/mrm.1910330323

Menon, R.S., Ogawa, S., Strupp, J.P., Uǧurbil, K., 1997. Ocular Dominance in Human V1 Demonstrated by Functional Magnetic Resonance Imaging. J Neurophysiol 77, 2780–2787. https://doi.org/10.1152/jn.1997.77.5.2780

Moeller, S., Pisharady, P.K., Ramanna, S., Lenglet, C., Wu, X., Dowdle, L., Yacoub, E., Uğurbil, K., Akçakaya, M., 2021. NOise reduction with DIstribution Corrected (NORDIC) PCA in dMRI with complex-valued parameter-free locally low-rank processing. Neuroimage 226, 117539–117539. https://doi.org/10.1016/j.neuroimage.2020.117539

O’Brien, K.R., Kober, T., Hagmann, P., Maeder, P., Marques, J., Lazeyras, F., Krueger, G., Roche, A., 2014. Robust T1-Weighted Structural Brain Imaging and Morphometry at 7T Using MP2RAGE. Plos One 9, e99676. https://doi.org/10.1371/journal.pone.0099676

Ogawa, S., Lee, T., Nayak, A.S., Glynn, P., 1990. Oxygenation-sensitive contrast in magnetic resonance image of rodent brain at high magnetic fields. Magnetic Resonance in Medicine. 14, 68–78. https://doi.org/10.1002/mrm.1910140108

Olman, C.A., Harel, N., Feinberg, D.A., He, S., Zhang, P., Ugurbil, K., Yacoub, E., 2012. Layer-specific fMRI reflects different neuronal computations at different depths in human V1. PloS one. 7, e32536. https://doi.org/10.1371/journal.pone.0032536

Peirce, J., Gray, J.R., Simpson, S., MacAskill, M., Höchenberger, R., Sogo, H., Kastman, E., Lindeløv, J.K., 2019. PsychoPy2: Experiments in behavior made easy. Behav Res Methods 51, 195–203. https://doi.org/10.3758/s13428-018-01193-y

Polimeni, and D.F.T.D., Alexander Beckett, An Thanh Vu, Jonathan, 2018. Blood-volume imaging using GRASE-VASO at ultra-high field for layer specific fMRI in human brain, in: Joint Annual Meeting ISMRM-ESMRMB 2018. https://doi.org/https://cds.ismrm.org/protected/18MProceedings/PDFfiles/0709.html

Polimeni, J.R., Fischl, B., Greve, D.N., Wald, L.L., 2010. Laminar analysis of 7 T BOLD using an imposed spatial activation pattern in human V1. Neuroimage. 52, 1334–1346. https://doi.org/10.1016/j.neuroimage.2010.05.005

Poser, B.A., Koopmans, P.J., Witzel, T., Wald, L.L., Barth, M., 2010. Three dimensional echo-planar imaging at 7 Tesla. Neuroimage. 51, 261–266. https://doi.org/10.1016/j.neuroimage.2010.01.108

Rockland, K.S., Pandya, D.N., 1979. Laminar origins and terminations of cortical connections of the occipital lobe in the rhesus monkey. Brain Research. 179, 3–20. https://doi.org/10.1016/0006-8993(79)90485-2

Salin, P.A., Bullier, J., 1995. Corticocortical connections in the visual system: structure and function. Physiol Rev 75, 107–154. https://doi.org/10.1152/physrev.1995.75.1.107

Self, M.W., Kerkoerle, T. van, Goebel, R., Roelfsema, P.R., 2017. Benchmarking laminar fMRI: neuronal spiking and synaptic activity during top-down and bottom-up processing in the different layers of cortex. Neuroimage 197, 806–817. https://doi.org/10.1016/j.neuroimage.2017.06.045

Stanley, O.W., Kuurstra, A.B., Klassen, L.M., Menon, R.S., Gati, J.S., 2021. Effects of phase regression on high-resolution functional MRI of the primary visual cortex. Neuroimage 227, 117631. https://doi.org/10.1016/j.neuroimage.2020.117631

Stirnberg, R., Stöcker, T., 2021. Segmented K-space blipped-controlled aliasing in parallel imaging for high spatiotemporal resolution EPI. Magnet Reson Med 85, 1540–1551. https://doi.org/10.1002/mrm.28486

Turner, R., 2002. How much cortex can a vein drain? Downstream dilution of activation-related cerebral blood oxygenation changes. Neuroimage. 16, 1062–1067. https://doi.org/10.1006/nimg.2002.1082

Uludağ, K., Blinder, P., 2018. Linking brain vascular physiology to hemodynamic response in ultra-high field MRI. Neuroimage. 168, 279–295. https://doi.org/10.1016/j.neuroimage.2017.02.063

Uludağ, K., Müller-Bierl, B., Uğurbil, K., 2009. An integrative model for neuronal activity-induced signal changes for gradient and spin echo functional imaging. Neuroimage. 48, 150–165. https://doi.org/10.1016/j.neuroimage.2009.05.051

Vizioli, L., Moeller, S., Dowdle, L., Akçakaya, M., Martino, F.D., Yacoub, E., Uğurbil, K., 2021. Lowering the thermal noise barrier in functional brain mapping with magnetic resonance imaging. Nat Commun 12, 5181. https://doi.org/10.1038/s41467-021-25431-8

Weber, B., Keller, A.L., Reichold, J., Logothetis, N.K., 2008a. The Microvascular System of the Striate and Extrastriate Visual Cortex of the Macaque. Cereb Cortex 18, 2318–2330. https://doi.org/10.1093/cercor/bhm259

Weber, B., Keller, A.L., Reichold, J., Logothetis, N.K., 2008b. The microvascular system of the striate and extrastriate visual cortex of the macaque. Cerebral Cortex. 18, 2318–2330. https://doi.org/10.1093/cercor/bhm259

Yacoub, E., Duong, T.Q., Moortele, P.V.D., Lindquist, M., Adriany, G., Kim, S., Uğurbil, K., Hu, X., 2003. Spin-echo fMRI in humans using high spatial resolutions and high magnetic fields. Magn. Reson. Med. 49, 655–664. https://doi.org/10.1002/mrm.10433

Yacoub, E., Shmuel, A., Logothetis, N., Uğurbil, K., 2007. Robust detection of ocular dominance columns in humans using Hahn Spin Echo BOLD functional MRI at 7 Tesla. Neuroimage 37, 1161–1177. https://doi.org/10.1016/j.neuroimage.2007.05.020

Zaretskaya, N., Bause, J., Polimeni, J.R., Grassi, P.R., Scheffler, K., Bartels, A., 2020. Eye-selective fMRI activity in human primary visual cortex: Comparison between 3 T and 9.4 T, and effects across cortical depth. Neuroimage 220, 117078. https://doi.org/10.1016/j.neuroimage.2020.117078

Zhao, F., Wang, P., Hendrich, K., Ugurbil, K., Kim, S.-G., 2006. Cortical layer-dependent BOLD and CBV responses measured by spin-echo and gradient-echo fMRI: insights into hemodynamic regulation. Neuroimage 30, 1149–1160. https://doi.org/https://doi.org/10.1016/j.neuroimage.2005.11.013

