## supplementary materials for "Layer Dependence of Monocular and Binocular Responses in Human Ocular Dominance Columns at 7T using VASO and BOLD"

### 1. Appendix A. Supplementary data

#### 1.1. BOLD and VASO laminar profiles

Figure S1 shows the BOLD and VASO statistical maps corresponding to both eyes, right eye, and left eye stimulations along with their laminar profiles after thermal noise removal using NORDIC [1,2]. Note the higher signal change in both contrasts after NORDIC which is visible in both the statistical t-maps and laminar profiles. Also, there is a slight difference between the peak location in monocular and binocular VASO responses *after* and *before* applying NORDIC. Before NORDIC, laminar VASO profiles peak at deep and middle depths, but after NORDIC the peak is mainly located at middle cortical depth.

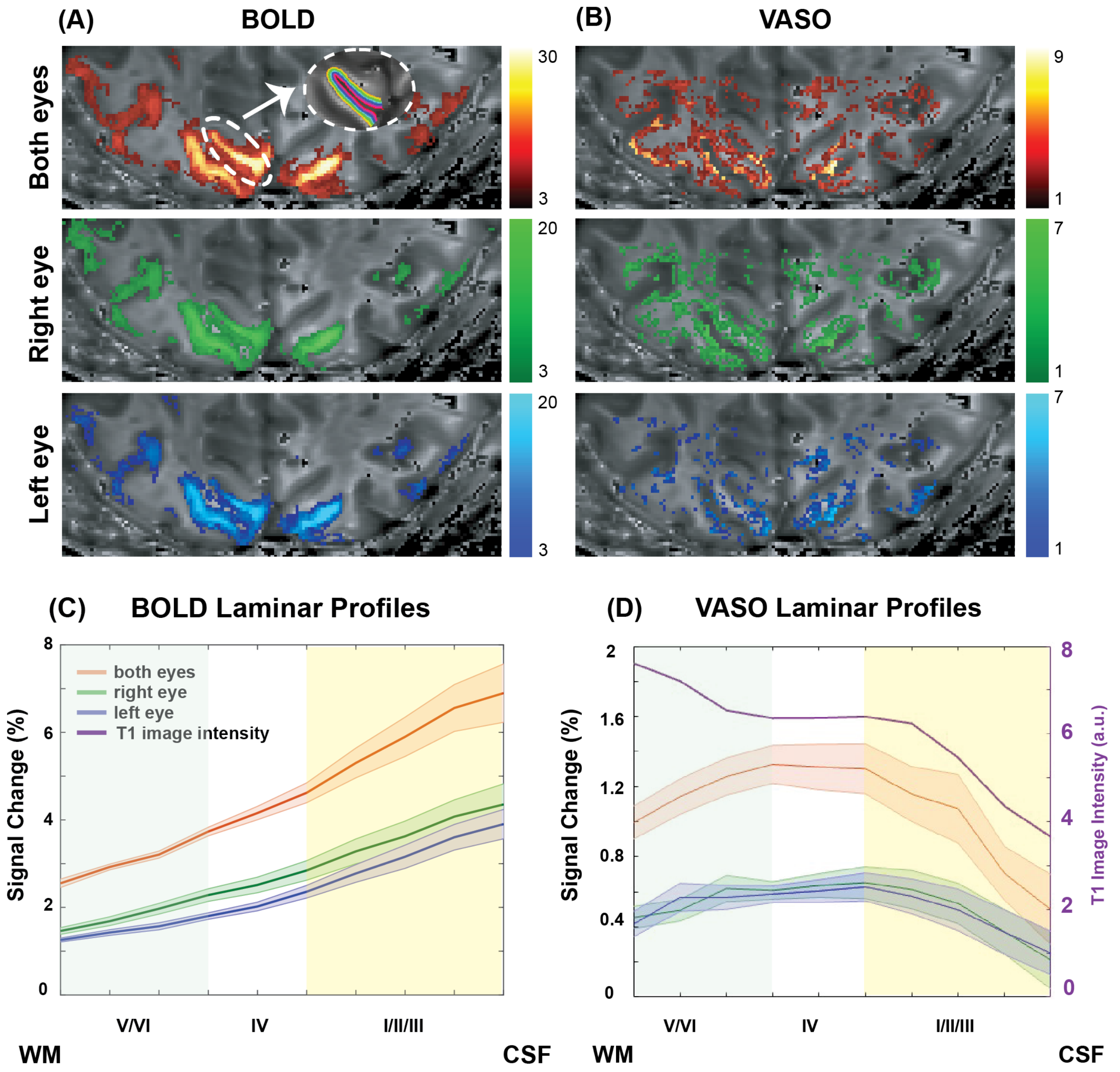


*Figure S1 : BOLD (A) and VASO (B) statistical maps (t-maps) overlaid on T1-EPI image of one representative participant. Laminar BOLD (C) and VASO (D) profiles corresponding to both eyes, right eye, and left eye stimulations.*

#### 1.2. Right and left eye responses to a monocular viewing

Figure S2 depicts the response of the right (left)-eye-dominated-columns to the right and left eye stimulation of the BOLD and VASO contrasts after applying NORDIC. The most noticeable difference between this figure and Figure 4 of the manuscript (same result but without NPRDIC) is the improved differentiability between the left-eye-dominated columns response to the “corresponding” and “opposite” eye stimulation. In addition, as mentioned earlier, the laminar VASO profiles peak at middle cortical depth after applying NORDIC, while the native profiles peak at deep and middle cortical depth.

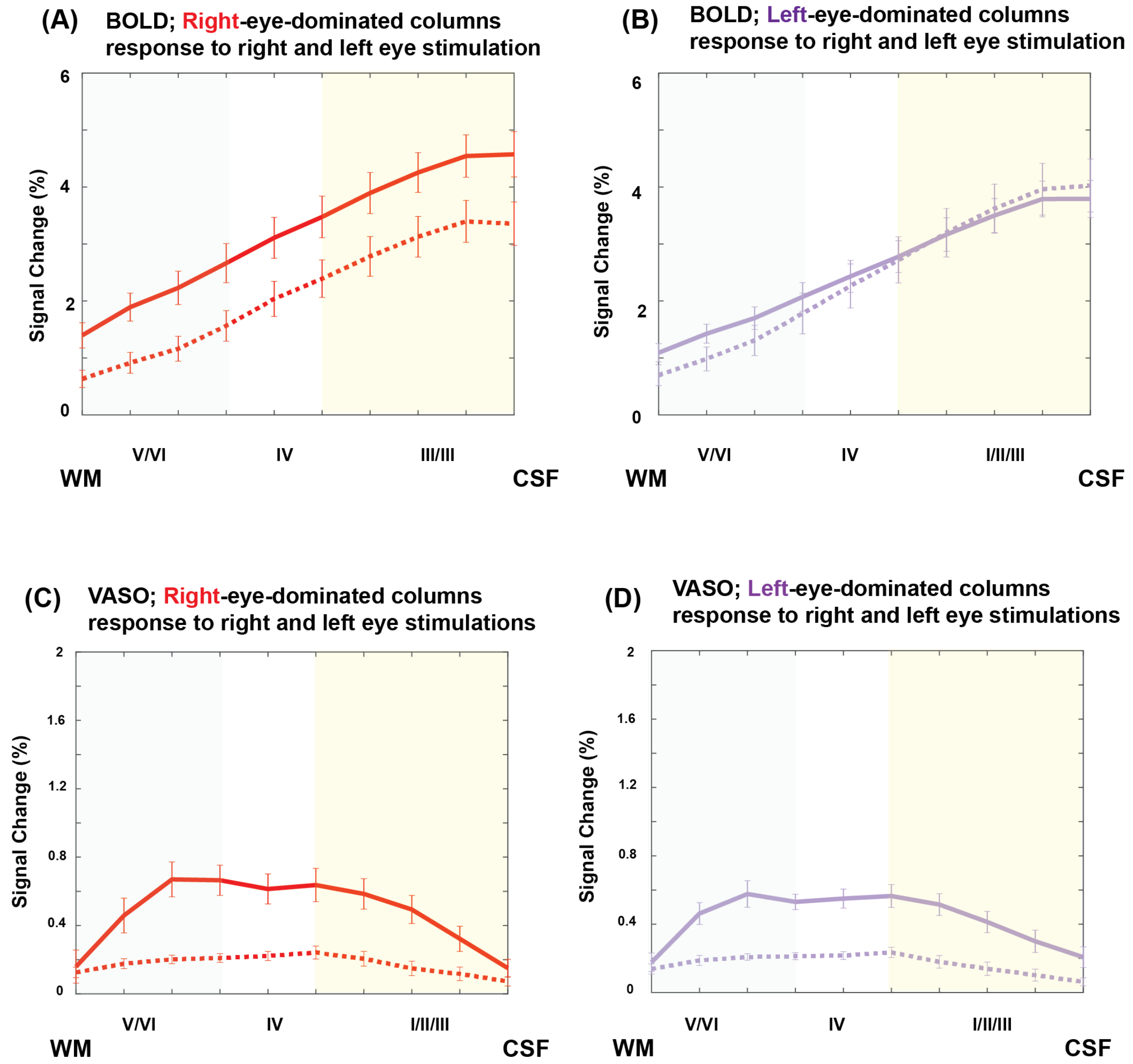


*Figure S2: The response of the right-eye-dominated-columns to the right and left eye stimulation in BOLD (A) and VASO (C) after thermal noise removal using NORDIC. Similarly, the response of the left-eye-dominated columns to the left and right stimulation in BOLD (B) and VASO (D). Error bars show the standard error of mean.*

#### 1.3. The response of the monocular columns to binocular viewing

Figure S3 illustrates the response of the monocular columns to a binocular stimulation after applying NORDIC. Similar to the results shown in Figures S1 and S2, both BOLD and VASO signal changes showed an increase in their amplitude after applying NORDIC. Consistent with the result shown in Figure 7 of the manuscript (same results but without NORDIC), the monocular columns response to the same eye and both eye stimulation was comparable, though with a slight reduction in deeper cortical depth. Running a paired t-test yielded a significant difference between the monocular columns responses to the "same eye" and "binocular stimulation" at *deep cortical depths* with $p = 0.001$ for the right eye responses and $p = 0.003$ for the left eye responses.

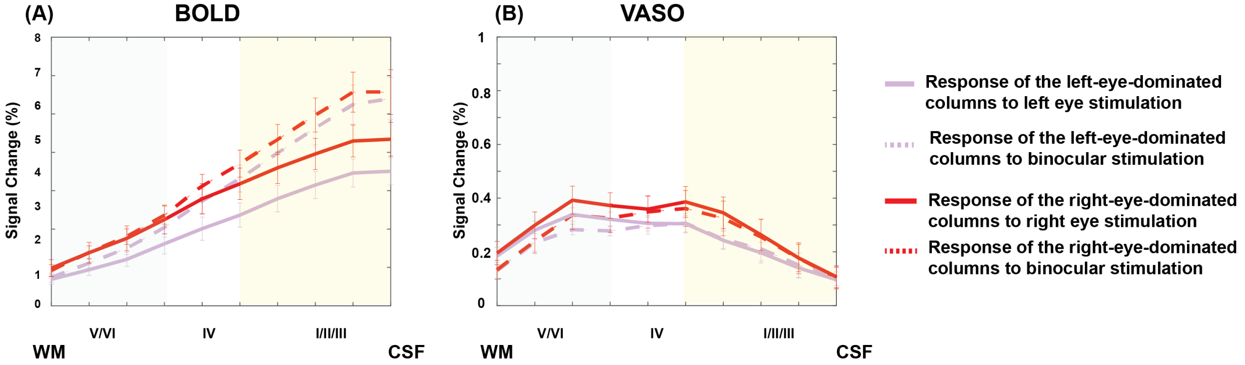


*Figure S3: The responses of monocular columns (i.e., right-eye-dominated columns and left-eye-dominated-columns) to monocular and binocular viewing in BOLD (A) and VASO (B) after thermal noise removal with NORDIC. Error bars show the standard error of the mean.*

Overall, NORDIC improved the image temporal signal-to-noise-ratio (tSNR) by the factor of ~ 1.5-2 and increased the laminar signal amplitude of both contrasts considerably. For instance, the VASO mean signal change across cortical depths associated with "both eyes stimulation" before NORDOC was 0.18 % and increased to 1.1 % after thermal noise removal with NORDIC. For BOLD, 3.1 % mean signal change across cortical depths correspond to "both eyes" stimulation increased to 4.6 % after applying NORDIC.

Almost all VASO profiles showed a hint of a double peak distribution across the cortical depths after applying NORDIC, which were not visible in any of these figures before applying NORDIC. Also note that at the single subject level, this double peak pattern was only visible in the dominant eye (right eye) responses.

Another interesting difference was improving the ocular segregation in VASO responses after thermal noise correction, especially for the left eye responses (compare Figures S2 C and S2 D with Figure 4C and 4D of the manuscript).

Please note that we performed NORDIC denoising with the default parameter settings (ARG.factor_error (FE) = 1).

**References:**

1. Moeller S, Pisharady PK, Ramanna S, Lenglet C, Wu X, Dowdle L, et al. NOise reduction with DIstribution Corrected (NORDIC) PCA in dMRI with complex-valued parameter-free locally low-rank processing. Neuroimage. 2021;226:117539–117539.

2. Vizioli L, Moeller S, Dowdle L, Akçakaya M, Martino FD, Yacoub E, et al. Lowering the thermal noise barrier in functional brain mapping with magnetic resonance imaging. Nat Commun. 2021;12:5181.
